# Members of the abscisic acid co-receptor PP2C protein family mediate salicylic acid-abscisic acid crosstalk

**DOI:** 10.1101/123059

**Authors:** Murli Manohar, Dekai Wang, Patricia M. Manosalva, Hyong Woo Choi, Erich Kombrink, Daniel F. Klessig

## Abstract

The interplay between abscisic acid (ABA) and salicylic acid (SA) influences plant responses to various (a)biotic stresses; however, the underlying mechanism(s) for this crosstalk is largely unknown. Here we report that type 2C protein phosphatases (PP2Cs), some of which are negative regulators of ABA signaling, bind SA. SA binding suppressed the ABA-enhanced interaction between these PP2Cs and various ABA receptors belonging to the PYR/PYL/RCAR protein family. Additionally, SA suppressed ABA-enhanced degradation of PP2Cs and ABA-induced stabilization of SnRK2s. Supporting SA’s role as a negative regulator of ABA signaling, exogenous SA suppressed ABA-induced gene expression, whereas SA-deficient *sid2-1* mutants displayed heightened PP2C degradation and hypersensitivity to ABA-induced suppression of seed germination. Together, these results suggest a new molecular mechanism through which SA antagonizes ABA signaling. A better understanding of the crosstalk between these hormones is important for improving the sustainability of agriculture in the face of climate change.

## Introduction

Elaborate hormone signaling networks allow plants to perceive and respond adaptively to various biotic and abiotic stresses (Tuteja, 2007; Raghavendra *et al*., 2010). One of the vital hormones that plays a central role in the adaptation to abiotic stresses, particularly drought and salt stresses, is ABA. In addition, ABA is involved in regulating plant growth and developmental processes under non-stress conditions (Raghavendra *et al*., 2010) and modulating defense responses following pathogen attack (Robert-Seilaniantz *et al*., 2011; Denance *et al*., 2013). Because of its essential role in multiple physiological processes both under stressed and non-stressed conditions, the ABA signaling pathway has been studied intensively during the last two decades. Initial attempts to identify ABA receptors were met with controversy and frustration. Several proteins were proposed to be ABA receptors, but their exact role in ABA response and their associated mechanisms were never established (Hauser *et al*., 2011).

The discovery that members of the pyrabactin resistance 1/PYR1-like/regulatory component of ABA receptor (PYR/PYL/RCAR) protein family are ABA receptors, and that they interact with members of the type 2C protein phosphatase (PP2C) protein subfamily, was a major breakthrough in dissecting the ABA signaling pathway (Fujii *et al*., 2009; Ma *et al*., 2009; Miyazono *et al*., 2009; Park *et al*., 2009; Santiago *et al*., 2009; Soon *et al*., 2012). In the absence of ABA, PP2Cs are able to bind and dephosphorylate members of the sucrose non-fermenting 1-related subfamily 2 protein kinase (SnRK2) family. This negatively regulates ABA signaling because autophosphorylation is required for SnRK2 kinase activity, and thus their ability to transduce the ABA signal by phosphorylating downstream targets. In the presence of ABA, the ABA-receptor complex tightly binds to PP2Cs, thereby preventing PP2C-mediated dephosphorylation of SnRK2. This, in turn, allows activated SnRK2s to relay the ABA signal.

The reversible phosphorylation of proteins by protein kinases and phosphatases is an important mechanism for regulating many biological processes. In contrast to eukaryotic protein kinases, whose primary and three-dimensional structures are very similar, protein phosphatases are diverse. Depending on their substrate specificity, protein phosphatases can be divided into two classes, serine/threonine (Ser/Thr) or tyrosine phosphatases (Schweighofer *et al*., 2004; Fuchs *et al*., 2013; Singh *et al*., 2015). The Ser/Thr phosphatases have been further organized into the phosphoprotein phosphatase (PPP) and metal-dependent protein phosphatase (PPM) families. In plants, PP2Cs, which belong to the PPM family, represent a major portion of the phosphatase-encoding gene family. To date, 80 or more genes have been identified in the Arabidopsis, tomato, rice and hot pepper genomes. Phylogenetic analyses have further divided the PP2C families from these plant species into ten or more subclades designated alphabetically from A onward (Fuchs *et al*., 2013; Singh *et al*., 2015).

Of the PP2C subclades, members of “clade A” have been studied the most extensively, as they negatively regulate ABA signaling in various plant species. In Arabidopsis, clade A proteins such as ABA-insensitive 1 (ABI1), ABI2, Hypersensitive to ABA 1 (HAB1), and PP2CA/AHG3, have been shown to mediate ABA-induced responses to abiotic and biotic stresses via their interaction with SnRK2s and PYR/PYL/RCARs (de Torres-Zabala *et al*., 2007; Fujii *et al*., 2009; Santiago *et al*., 2012; Soon *et al*., 2012; Lim *et al*., 2014). Functional studies of PP2C proteins from other clades are limited, but they suggest that some of these proteins are involved in responding to (a)biotic stresses. For instance, the clade B member AP2C1 (*Arabidopsis* phosphatase 2C1) and its ortholog MP2C from *Medicago sativa* regulate the activity of stress-induced mitogen-activated protein kinases (MAPKs; (Meskiene *et al*., 2003; Schweighofer *et al*., 2004), and the clade F member PIA1 (PP2C induced by AvrRpm1) regulates immune responses in Arabidopsis (Widjaja *et al*., 2010). By contrast, clades C and D contain PP2Cs that regulate developmental processes (Schweighofer *et al*., 2004; Fuchs *et al*., 2013; Singh *et al*., 2015). Members of clade C, including POL (Poltergeist) and PLL (POL-like), control shoot and root meristem formation and embryo formation (Song & Clark, 2005), whereas members of clade D negatively regulate the activity of plasma membrane H^+^-ATPases, and thus cell expansion in the absence of auxin (Spartz *et al*., 2014).

SA is another important plant hormone involved in diverse physiological and metabolic processes, including plant responses to various abiotic stresses. In addition, SA is an essential regulator of plant immune responses (Vlot *et al*., 2009; Manohar *et al*., 2015; Klessig *et al*., 2016). While several recent studies have identified components of SA signaling networks and revealed some SA-mediated signaling mechanisms, a full picture of SA-based signaling in plants is far from complete. Indeed, the identity of the SA receptor(s) remains unclear. It was recently proposed that Non-expresser of PR1 (NPR1), which functions as a master regulator of SA-mediated immune signaling, is an SA receptor (Wu *et al*., 2012). In contrast, Fu and coworkers (2012) suggested that NPRl’s two homologs, NPR3 and NPR4, rather than NPR1, are SA receptors. Since NPR3 and NPR4 are adaptors for Cullin 3 ubiquitin E3 ligase, they may regulate the SA signaling pathway by fine-tuning NPR1 protein levels via degradation (Fu *et al*., 2012). In addition, nearly 30 SA-binding proteins (SABPs) have been identified (Klessig et al., 2016). These proteins exhibit a wide range of affinities for SA, and SA binding alters their activities. Given that SA levels vary dramatically within a plant depending on the subcellular compartment, tissue type, developmental stage and external cues, such as infection, these findings raise the possibility that SA exerts its effects by interacting with multiple targets, rather than a small number of receptors.

Although SA’s role in activating disease resistance and ABA’s role in signaling abiotic stress responses are well-known, it is only recently becoming apparent that ABA also influences immune responses (Robert-Seilaniantz *et al*., 2011; Denance *et al*., 2013). ABA treatment suppressed defense responses and enhanced plant susceptibility to certain bacterial and fungal pathogens (Ward *et al*., 1989; McDonald & Cahill, 1999; Mohr & Cahill, 2003; Thaler & Bostock, 2004; De Torres Zabala *et al*., 2009; Robert-Seilaniantz *et al*., 2011). Additionally, the virulence of *Pseudomonas syringae* in Arabidopsis was dependent on manipulation of the ABA signaling pathway by secreted bacterial effectors (de Torres-Zabala *et al*., 2007). Growing evidence also indicates that there is substantial crosstalk between the ABA and SA pathways during immune signaling (de Torres-Zabala *et al*., 2007; Yasuda *et al*., 2008). Arabidopsis mutants deficient in ABA synthesis or response not only exhibited reduced susceptibility to pathogen infection, but also showed enhanced expression of SA-responsive genes, such as *Pathogenesis-Related Protein-1* (*PR-1*) and *PR-4* (Audenaert *et al*., 2002; Thaler & Bostock, 2004; Sanchez-Vallet *et al*., 2012). Conversely, Arabidopsis overexpressing RCAR3, which confers increased ABA sensitivity, displayed enhanced susceptibility to *P. syringae* DC3000 infection, which correlated with decreased expression of *PR-1* and *NPR1* (Lim *et al*., 2014). Further demonstrating the antagonistic interaction between ABA and SA, exogenous ABA suppressed the ability of an SA functional analog to enhance pathogen resistance in Arabidopsis, while pretreatment with this analog suppressed NaCl-induced expression of several ABA biosynthetic or responsive genes (Yasuda *et al*., 2008). ABA appears to suppress immune responses by down-regulating SA biosynthesis (de Torres-Zabala *et al*., 2007; Yasuda *et al*., 2008); however, the mechanism through which SA inhibits ABA signaling is unknown.

In previous studies, we have identified several SABPs that are involved in various biotic and abiotic stress responses (Tian *et al*., 2012; Manohar *et al*., 2015). Here, we identify PP2Cs from clade A and D as novel SABPs and show that SA binding to these PP2Cs suppresses their ABA-enhanced interaction with the ABA receptors. In addition, SA suppresses ABA-induced degradation of PP2Cs and suppresses ABA-mediated stabilization of SnRK2s. Combined with the demonstration that SA treatment antagonizes ABA-induced gene expression and SA deficient *sid2-1* mutants are ABA hypersensitive, these results suggest that SA antagonizes ABA signaling through multiple mechanisms.

## Results

### Identification of PP2Cs as novel SA-binding proteins

To help define the SA signaling network in plants, we developed several high-throughput screens capable of identifying SABPs on a genome-wide scale (Tian *et al*., 2012; Choi *et al*., 2015; Manohar *et al*., 2015). In one screen, protein extracts prepared from Arabidopsis leaves were subjected to affinity chromatography on a Pharmalink column to which SA was attached. After stringent washing with the biologically inactive SA analog 4-hydroxy benzoic acid (4-HBA), SA-bound proteins on the column were eluted with excess SA. The eluted proteins were analyzed by mass spectroscopy and a PP2C belonging to clade D (PP2C-D4; At3g55050) was identified along with other putative SABPs. Recombinant histidine-tagged PP2C-D4 was produced in *Escherichia coli* and the purified protein was further assessed for SA-binding activity using three different assays, including surface plasmon resonance (SPR), photoaffinity crosslinking, and size-exclusion chromatography. SPR analysis was performed with a CM5 sensor chip to which the SA derivative 3-aminoethyl SA (3AESA) was immobilized via an amide bond. Binding to 3AESA was detected when purified PP2C-D4 was passed over the sensor chip (Figure 1A). In the presence of increasing concentrations of SA, PP2C-D4 binding to the 3AESA-immobilized sensor chip was modestly reduced (Figure 1A). Similar to these results, the photoaffinity labeling approach indicated that PP2C-D4 bound and was crosslinked to 4-azido SA (4AzSA). This binding also was suppressed by SA in a dose-dependent manner (Figure 1B), arguing that PP2C-D4 binding to both 3AESA and 4AzSA represents authentic SA-binding activity. PP2C-D4’s ability to bind SA also was confirmed by size-exclusion chromatography using [^3^H]SA; binding to [^3^H]SA was partially suppressed by excess unlabeled SA, but not by excess amount of 4-HBA (Figure 1C). An independent, parallel screen using a SA-derived ligand in combination with the yeast three-hybrid technology, which relies on the *in vivo* interaction between the ligand (small molecule) and its protein target in the yeast nucleus, also identified PP2C-D4 (called PP2C6 in (Cottier *et al*., 2011)). Based on these five independent assays, we conclude that PP2C-D4 is a true SABP.

**Figure 1:**
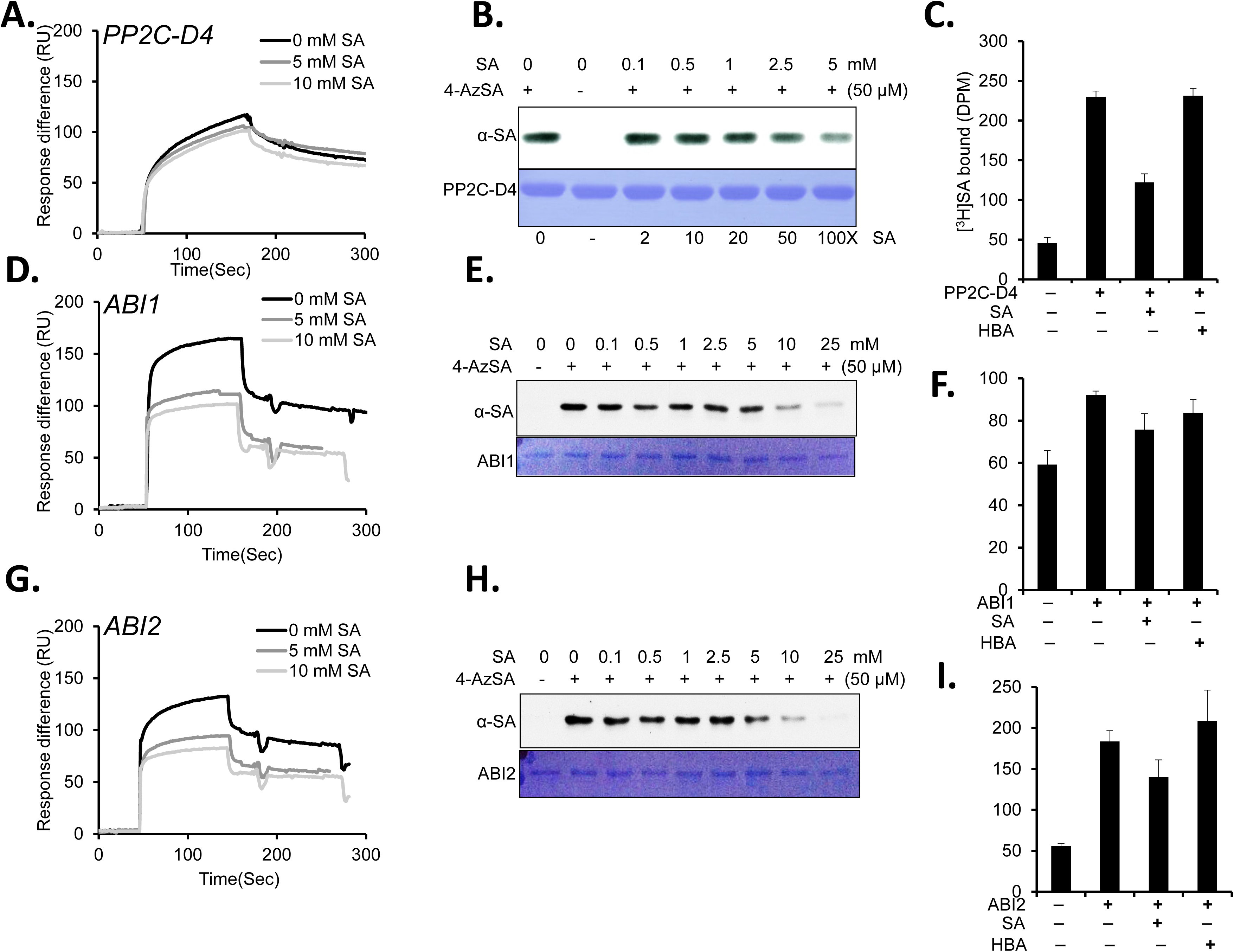
Several PP2Cs bind SA. **(A, D, G)** Sensorgrams for 1 μΜ of His_6_-tagged PP2C-D4 (At3g55050) **(A)**, ABI1 (At4g26080) **(D)**, and ABI2 (At5g57050) **(G)** in the absence (0 mM) or presence of two concentrations of SA (5 or 10 mM) using a 3AESA-immobilized SPR sensor chip. Signals detected from a mock-coupled control chip were subtracted. **(B, E, H)** Photo-activated crosslinking of 50 ng of PP2C-D4 **(B)**, ABI1 **(E)**, and ABI2 **(H)** to 4AzSA (50 μΜ) in the absence or presence of increasing amounts of SA was detected by immunoblotting using an α-SA antibody. Reactions without 4AzSA served as negative controls. Proteins stained with Coomassie Brilliant Blue (CBB) served as the loading controls. **(C, F, I)** Binding of [^3^H]SA (200 nM) by 200 ng/μl PP2C-D4 **(C)**, ABI1 **(F)**, and ABI2 **(I)** in the absence or presence of a 10,000-fold excess of unlabeled SA was determined by size-exclusion chromatography. Chromatography with [^3^H]SA in the absence of protein served as negative controls. Reactions with [^3^H]SA with excess of 4-amino benzoic acid (4-HBA), an inactive SA analog, served as negative controls for SA-specific competitive inhibition. The experiments was independently repeated at least twice.

Several clade A PP2C family members, including ABI1, ABI2, and HAB1, have been identified as core components of the ABA signaling network (Soon *et al*., 2012). This prompted us to test whether these proteins also bind SA. Like PP2C-D4, recombinant ABI1 and ABI2 bound the 3AESA-immobilized sensor chip and crosslinked with 4AzSA (Figure 1D, 1E, 1G, 1H). The ability of ABI1 and ABI2 to bind 3AESA and crosslink to 4AzSA also was partially suppressed by SA in a dose-dependent manner. ABI2’s binding to [^3^H]SA was comparable to that of PP2C-D4 and this binding was partially suppressed by excess unlabeled SA but not by excess 4-HBA, while ABI1 exhibited relatively weak binding to [^3^H]SA and suppression by excess unlabeled SA (Figure 1F; 1I). Interestingly, SA suppressed the binding of these proteins to 3AESA more effectively than that of PP2C-D4 (Figure 1D, 1G). In contrast, HAB1 displayed much weaker binding to the 3AESA-immobilized sensor chip (Supporting Figure 1A). The ability of phosphoprotein phosphatase 2A regulatory subunit A (PP2A), a component of phosphatases belonging to the PPP family, also was tested for SA binding. This protein was previously identified during our screens, but it failed to meet the criteria as an SABP (Manohar *et al*., 2015). Consistent with these results, PP2A exhibited very weak binding to the 3AESA-immobilized sensor chip (Supporting Figure 1B). Whether other key components involved in ABA signaling also bind SA was then assessed. Size-exclusion chromatography revealed little or no binding of [^3^H]SA by three members of the PYR/PYL/RCAR family of ABA receptors (PYL1, PYL2, and PYR1) or by three SnRK2s (SnRK2.2, 2.3, and 2.6; Supporting Figure 1C). Likewise, SnRK2.2 displayed only very low-level binding to the 3AESA-immobilized sensor chip (Supporting Figure 1D). Together, these results suggest that SA preferentially interacts with specific PP2C family members, but not with other major components of the ABA signaling pathway.

Since clade A PP2Cs are negative regulators of the ABA signaling pathway, we tested whether the presence of ABA affects PP2C-SA interactions by flowing the PP2Cs over 3AESA-immobilized sensor chips in the absence or presence of ABA. Notably, binding of PP2C-D4, ABI1, and ABI2 to 3AESA was significantly enhanced in the presence of ABA (Figure 2). This ABA-induced enhancement was suppressed by excess SA, further arguing that 3AESA binding by these PP2Cs represents authentic SA-binding activity. In contrast, ABA failed to enhance the weak binding of HAB1 and PP2A or to enable SnRK2.2 to bind 3AESA (Supporting Figure 1A, 1B, 1D).

**Figure 2:**
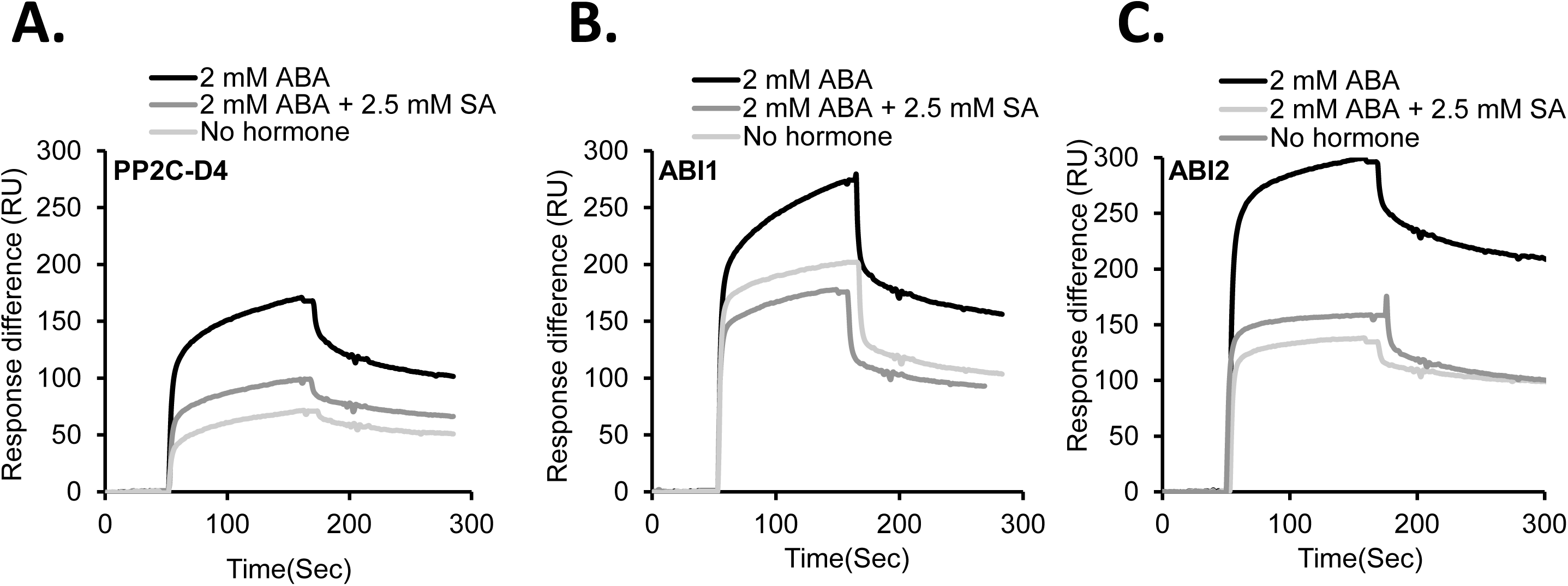
SA-binding activities of PP2Cs are enhanced by ABA and this binding is partially suppressed by SA. **(A-C)** Sensorgrams obtained with recombinant, purified 1 μΜ of His_6_-tagged PP2C-D4 **(A)**, ABI1 **(B)**, and ABI2 **(C)** using a 3AESA-immobilized sensor chip in the absence or in the presence of 2 mM ABA or 2 mM ABA plus 2.5 mM SA. Signals detected from a mock-coupled control chip were subtracted. The experiments was independently repeated at least twice.

### SA suppresses the ABA-enhanced interaction between PP2Cs and ABA receptors

To assess whether SA binding by PP2Cs alters their ability to interact with ABA receptors, SPR was performed. Purified PYL1 was immobilized on the CM5 sensor chip via an amide bond and interactions were detected by flowing purified PP2C-D4, ABI1, or ABI2 over the sensor chip.

Dose-dependent binding responses were obtained with all three PP2Cs tested, and this binding was enhanced in the presence of ABA (Figure 3; Supporting Figure 2). Notably, SA partially suppressed the ABA-enhanced interaction between PYL1 and all three PP2Cs (Figure 3). In the absence of ABA, SA slightly enhanced the interaction between PYL1 and PP2C-D4, but modestly suppressed binding by ABI1 and ABI2. SPR analysis with PYL2 and PYR1 revealed similarly dosage-dependent binding to the PP2Cs, that was further enhanced in the presence of ABA (Supporting Figures 3 & 4). Furthermore, this ABA-enhanced binding was suppressed by SA, although the level of suppression varied depending on the identity of the interacting proteins. For example, SA only weakly suppressed the ABA-enhanced interactions between PYR1 and all three PP2Cs, while interactions between these PP2Cs and either PYL1 or PYL2 were more strongly suppressed by SA (Figure 3, Supporting Figures 3 & 4). The ABA-enhanced interactions between PP2C-D4 and PYL1 or PYL2 also were suppressed less effectively by SA than the interactions between these ABA receptors and other PP2Cs (Figure 3, Supporting Figure 3). Interestingly, SA alone consistently enhanced the interaction between PP2C-D4 and all three ABA receptors, in some cases by a substantial amount (Figure 3, Supporting Figures 3 & 4).

**Figure 3:**
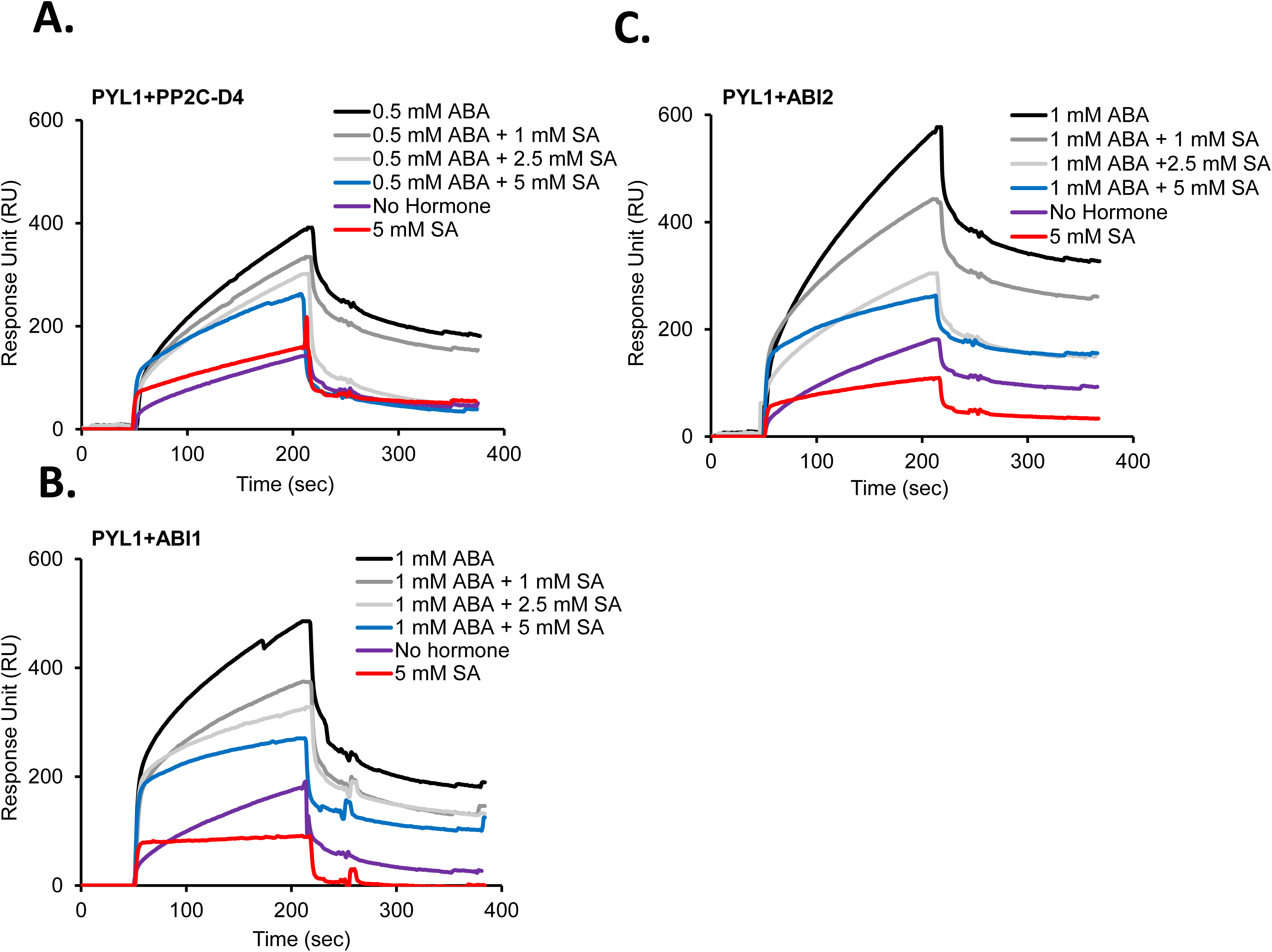
SA disrupts the ABA-induced interaction between PP2Cs and PYL1. **(A-C)** Sensorgrams obtained with recombinant, purified 10 μΜ of His_6_-tagged PP2C-D4 **(A)**, 1 μΜ of ABI1 **(B)** or 2.5 μM of ABI2 **(C)** using a His_6_-SUMO tagged PYL1-immobilized sensor chip in the absence or presence of the indicated concentrations of ABA or SA. Signals detected from a mock-coupled control chip were subtracted. The experiments was independently repeated at least twice.

### SA suppresses ABA-enhanced degradation of PP2Cs and ABA-induced stabilization of SnRK2s

Proteolysis plays an important role in regulating plant responses to various stresses by fine-tuning the turnover of key signaling components (Vierstra, 2009). Recent studies in Arabidopsis and rice have demonstrated that all three key components of ABA signaling, including PYR/PYL/RCARs, PP2Cs, and SnRK2s, are regulated by controlled proteolysis (Irigoyen *et al*., 2014; Kong *et al*., 2015; Lin *et al*., 2015). For example, ABA promotes degradation of ABI1, but it suppresses degradation of certain PYR/PYL/RCARs and SnRK2s (Kong *et al*., 2015; Lin *et al*., 2015). Furthermore, the plant hormone gibberellic acid (GA) antagonizes ABA signaling, in part, by stimulating degradation of PYR/PYL/RCARs and SnRK2s (Lin *et al*., 2015). To determine whether SA affects protein turnover, we analyze the stability of purified recombinant PP2Cs and SnRK2s in a cell-free degradation assay. Following incubation in protein extracts prepared from Arabidopsis seedlings supplemented with ABA and/or SA, immunoblot analyses indicated that the levels of His6-tagged PP2C-D4, ABI1, and ABI2 decreased in extracts supplemented with 10 μM ABA (Figure 4A). By contrast, the levels of these proteins remained fairly stable in extracts supplemented with both ABA and SA. SA alone had little effect on ABI1 or ABI2 levels but a modest decrease of PP2C-D4 levels was detected (Figure 4A). Together, these results suggest that ABA enhances PP2C degradation, and that this heightened turnover is suppressed by SA.

**Figure 4:**
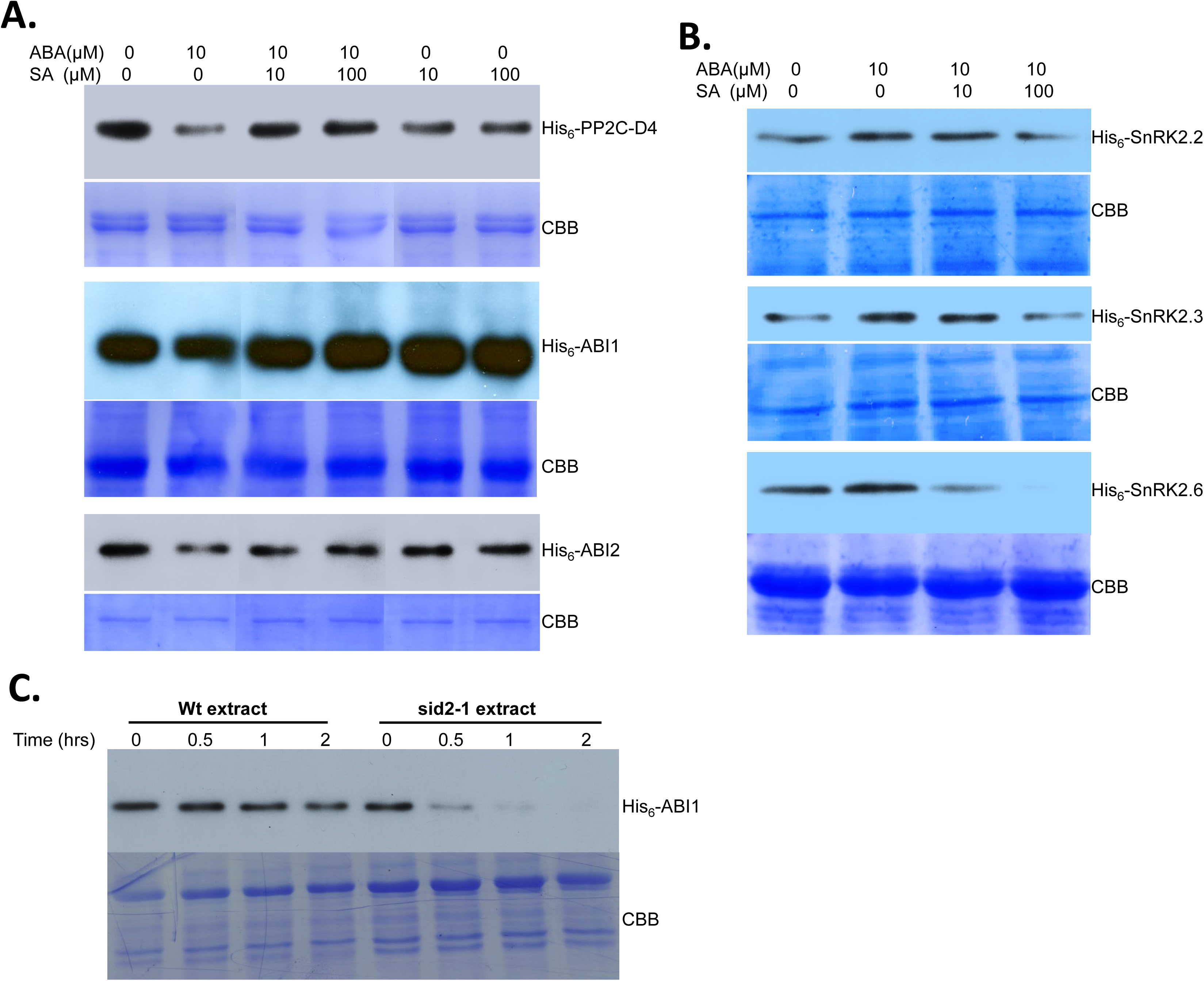
SA alters ABA-mediated turnover of PP2Cs and SnRK2s. **(A)** Cell-free degradation assay using total protein extracts prepared from ten-day-old Arabidopsis seedlings supplemented with 500 ng of His_6_-tagged PP2Cs (PP2C-D4, ABI1, and ABI2) and indicated concentrations of ABA, SA, or ABA+SA. **(B)** Cell-free degradation assay using total protein extracts prepared from ten-day-old Arabidopsis seedlings supplemented with 500 ng of His_6_-Sumo-tagged SnRK2s (SnRK2.2, 2.3, and 2.6) and indicated concentrations of ABA or ABA+SA. For A & B, the degradation assay was carried out at 30^°^ C for 3 hrs. All lanes shown are from the same experiment; some lanes unrelated to this study have been removed and lanes were then merged for clarity of presentation. **(C)** Cell-free degradation assay using total protein extracts prepared from ten-day-old wild-type or *sid2-1* Arabidopsis seedlings supplemented with 500 ng of His_6_-tagged ABI1. Samples were taken after 0, 0.5, 1, or 2 hrs of incubation; proteolysis was stopped by addition of SDS-PAGE buffer. Proteins were detected by immunoblotting using an α-His_6_-HRP antibody. Staining with Coomassie brilliant blue (CBB) staining of the gel served as a loading control. The experiments was independently repeated at least twice.

The stability of three SnRK2s, SnRK2.2, SnRK2.3, and SnRK2.6, was then assessed using the cell-free protein degradation assay. The levels of all three recombinant SnRK2s were slightly greater in extracts supplemented with 10 μM ABA as compared with unsupplemented extracts (Figure 4B). By contrast, SnRK2 levels were reduced in extracts containing both ABA and SA, with the greatest decrease detected after supplementation with ABA and 100 μM SA. Thus, ABA appears to stabilize SnRK2s, while SA suppresses ABA’s effect. Analysis of PYL1 did not reveal any change in protein levels regardless of supplementation with ABA and/or SA, suggesting that these hormones do not affect PYL1 stability (Supporting Figure 5).

The above results raised the possibility that endogenous SA antagonizes ABA signaling, at least in part, by stabilizing PP2Cs. To further assess this, the rate of ABI1 degradation was compared in protein extracts prepared from wild-type (WT) plants and the SA biosynthesis-deficient mutant *sid2-1*. ABI1 levels in the extract from *sid2-1* plants decreased substantially by 30 min and were barely detectable after 1 hour, whereas those in the WT extract decreased gradually over time (Figure 4C). Surprisingly, the enhanced degradation observed in *sid2-1* extracts was not reversed by i) adding SA to the extract, ii) spraying SA on *sid2-1* plants, or iii) supplementing *sid2-1* growth media with SA (Supporting Figure 6). Thus, while these results suggest that SA stabilizes ABI1, the failure of exogenous SA to slow ABI1 degradation in *sid2-1* extracts suggests that another factor(s) might be involved in this process.

### SA antagonizes ABA-induced gene expression *in vivo*

Previous studies have demonstrated that exogenously supplied ABA induces the accumulation of *ABI1* and *ABI2* transcripts (Hoth et al., 2002). To determine whether SA antagonizes the expression of these ABA signaling components, transcript levels for *ABI1* and *ABI2*, as well as *PP2C-D4*, were monitored in ABA- and/or SA- treated Arabidopsis. Quantitative reverse transcriptase-PCR (qRT-PCR) analyses showed that transcripts for *ABI1* and *ABI2* accumulated after ABA, but not SA, treatment (Figure 5A). An intermediate level of transcripts was detected in plants treated with SA and ABA, suggesting that the ABA-induced expression of these genes was partially suppressed by SA (Figure 5A). In comparison to the clade A PP2Cs, transcript accumulation for *PP2C-D4* was reduced in plants treated with either ABA or SA; an even greater reduction was observed in plants treated with both hormones. The expression of two well-known ABA-responsive genes, Response to Desiccation 29A (*RD29A*) and ABA-Responsive Element Binding Protein 2 (*AREB2*), also was analyzed. Consistent with previous studies, the expression of *RD29A* and *AREB2* was induced by ABA (Figure 5B) (Uno *et al*., 2000; Nakashima *et al*., 2006). Importantly, plants treated with ABA and SA accumulated reduced levels of *RD29A* and *AREB2* transcripts, indicating that the ABA-induced expression of these genes is partially suppressed by SA. By contrast, SA alone did not affect the expression of either gene.

**Figure 5:**
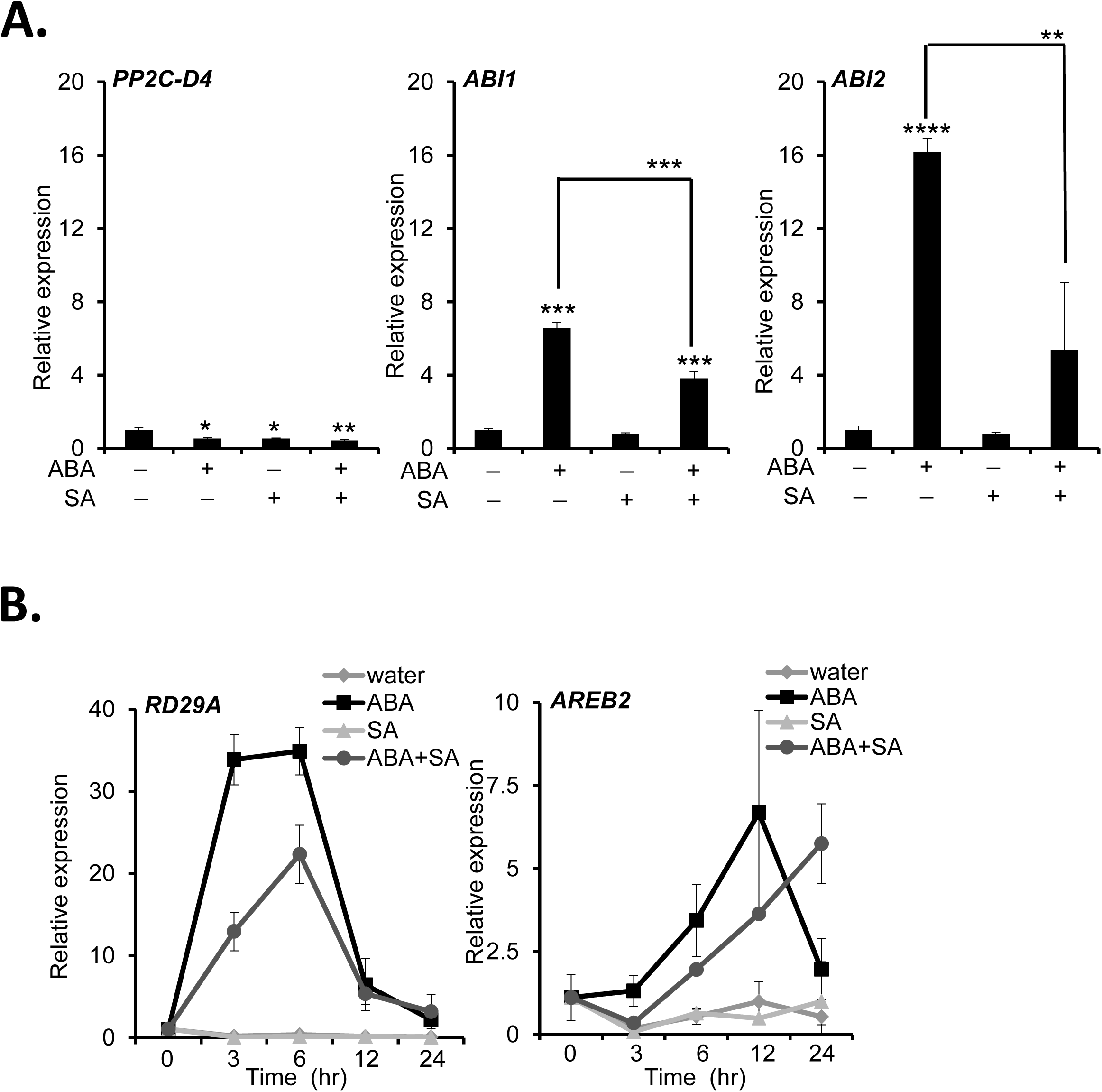
SA suppresses ABA-induced gene expression. **(A)** Transcript levels, as measured by qRT-PCR in seedlings pretreated with either water, 100 μΜ ABA, 100 μΜ SA or 100 μΜ ABA plus 100 μM SA. Transcript levels of *PP2C-D4, ABI1*, and *ABI2* were determined at 3 hrs post treatment (hpt). Data are averaged ± s.d (n=3). **(B)** Transcript levels as measured by qRT-PCR of ABA-responsive marker genes in seedlings pretreated with either water, 100 μΜ ABA, 100 μΜ SA or 100 μΜ ABA plus 100 μΜ SA. Transcript levels of *RD29A* (RESPONSIVE TO DESICCATION 29A) and *AREB2* (ABRE BINDING FACTOR 2) were determined at 0, 3, 6, 12 and 24 hpt. The relative expression levels were quantified by normalizing to ubiquitin expression level. Data are averaged ± s.d (n=4). *P ≤ 0.05; **P ≤ 0.005; ***P ≤ 0.0005; ****P ≤ 0.00005; two-tailed *t*-test. The experiments was independently repeated at least twice.

### An SA-deficient mutant is more sensitive to ABA-mediated seed dormancy

In addition to (a)biotic stress responses, ABA is involved in growth and developmental processes, including maintaining seed dormancy to prevent untimely germination (Kermode, 2005; Hubbard *et al*., 2010). To investigate whether SA antagonizes ABA’s ability to suppress germination, we monitored Arabidopsis seed germination on plates containing Murashige and Skoog (MS) medium in the presence or absence of ABA and/or SA. In the presence of 1 μΜ ABA, germination was dramatically reduced at all times monitored (Figure 6A). By contrast, 10 μΜ SA reduced germination slightly at 36 hrs, but from 48 hrs onward, the germination percentage of SA-treated and control seeds was comparable. Plates containing both SA and ABA displayed an intermediate level of germination. Thus, SA appears to suppress ABA-mediated inhibition of germination. Whether endogenous SA levels also affect ABA-mediated suppression of germination was then tested by comparing the germination of WT and *sid2-1* seeds. The germination rate for *sid2-1* seeds grown on ABA-containing plates was consistently lower than that of comparably grown WT seeds; by 72 hours, 15% of the *sid2-1* seeds had germinated in the presence of 1μΜ ABA, in contrast to 40% of WT seeds (Figure 6B). While SA completely overcame ABA suppression of seed germination in WT at 96 hrs post plating, it only partially reversed ABA’s effect in *sid2-1*. Based on the ABA hypersensitive phenotype displayed by *sid2-1* seeds, endogenous SA appears to play an important role in antagonizing ABA-mediated suppression of seed germination *in planta*.

**Figure 6:**
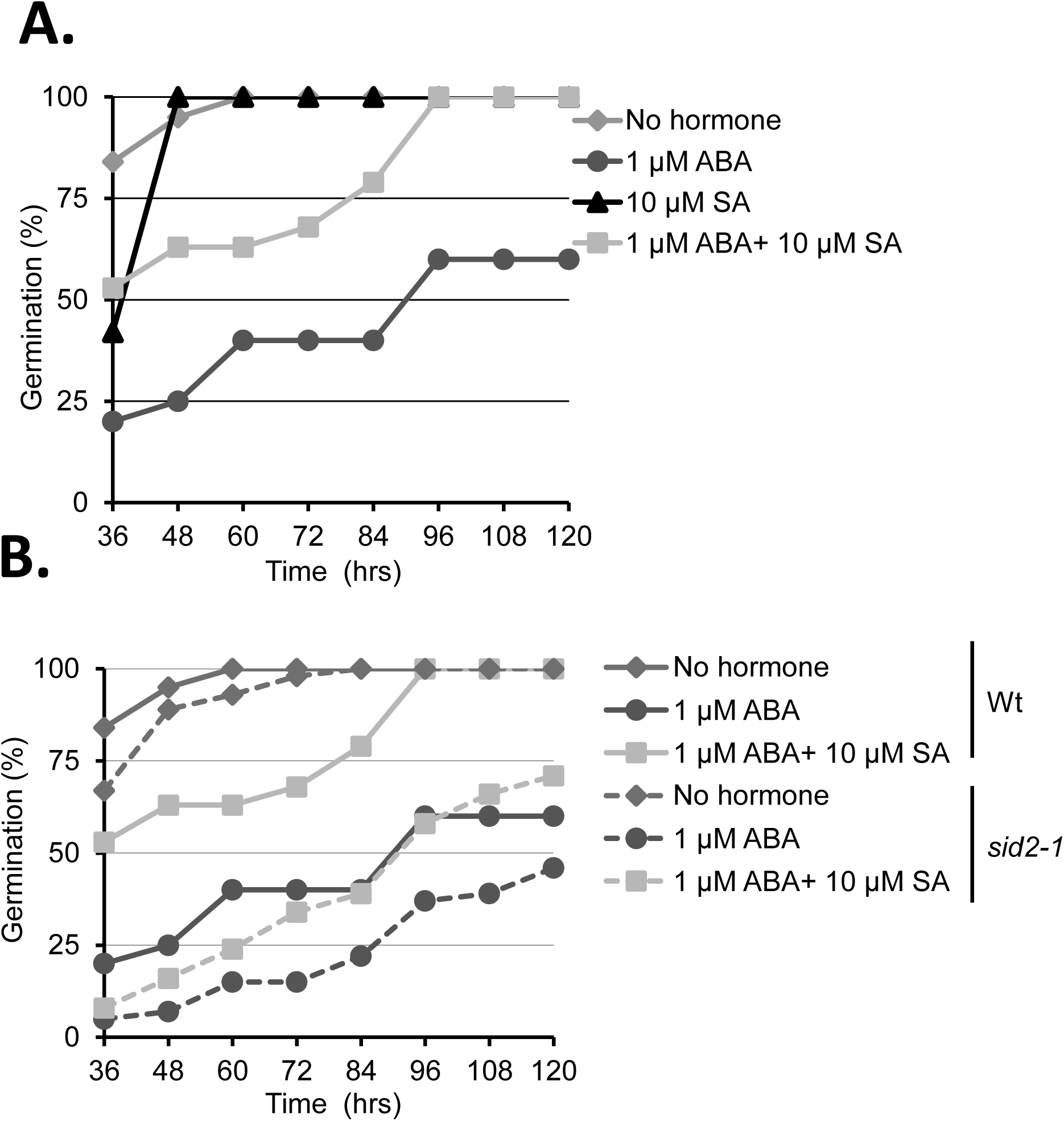
SA antagonizes ABA-mediated suppression of seed germination. **(A)** Germination rate of Arabidopsis wild-type seeds on MS medium containing no hormone or in the presence of 1 μΜ ABA, 10 μΜ SA or 1μΜ ABA plus 10 μΜ SA. **(B)** Comparison of germination rate between wild-type and *sid2-1* on MS medium containing no hormone or in the presence of 1 μΜ ABA or 1μΜ ABA plus 10 μΜ SA. The germination time course for wild-type seeds is shown with a solid line, while for *sid2-1* it is shown with a broken line (n=40 seeds). The percent of germinated seeds was determined at 36, 48, 60, 72, 84, 96, 108 and 120 hrs post plating. The experiments was independently repeated at least twice.

## Discussion

Elucidating the crosstalk between biotic and abiotic stress signaling pathways in plants is a rapidly expanding area of research. There is a growing recognition that ABA not only regulates abiotic stress responses and developmental processes, but also impacts plant-pathogen interactions (Robert-Seilaniantz *et al*., 2011; Denance *et al*., 2013). Likewise, SA not only signals plant immunity (Vlot *et al*., 2009; Manohar *et al*., 2015; Klessig *et al*., 2016), but also regulates responses to abiotic stresses and various aspects of growth and development (Hayat *et al*., 2010; Khan *et al*., 2015). To gain insights into how SA exerts its myriad effects, we previously developed several high throughput screens for identifying SABPs (Tian *et al*., 2012; Choi *et al*., 2015; Manohar *et al*., 2015). Here we report that several PP2Cs, including PP2C-D4, a member of clade D, and ABI1 and ABI2, members of clade A, are novel SABPs. By contrast, SA binding was not detected for two other phosphatases, HAB1 or PP2A, or for other components of the ABA signaling pathway, including various PYR/PYL/RCARs and SnRK2s. SPR analysis also revealed that binding of PP2C-D4, ABI1 and ABI2 to the SA analog 3AESA was enhanced in the presence of ABA. This finding suggests that PP2Cs from both clade A and D bind both ABA and SA, and that they do so in a cooperative manner. Although a previous report failed to detect binding between ABI1 and ABA (Ma *et al*., 2009), this discrepancy may be due to the very high sensitivity of SPR. Indeed, Ma et al. (2009) noted that ABI1 phosphatase activity was reduced up to 20% in presence of ABA.

Since clade A PP2Cs are critical negative regulators of ABA signaling, the discovery that they bind SA suggested that they play a role in modulating SA/ABA crosstalk. Several studies have documented an antagonistic relationship between SA and the ABA signaling pathway. For example, SA suppressed ABA-mediated inhibition of shoot growth and expression of cell cycle-related genes in rice (Meguro & Sato, 2014). Likewise, pretreating Arabidopsis with a compound that activates SA-dependent defense signaling antagonized the induction of ABA biosynthesis-related and ABA-responsive genes after NaCl treatment (Yasuda *et al*., 2008). Expanding on these findings, we demonstrated that SA treatment suppresses ABA-induced expression of the ABA signaling components *ABI1* and *ABI2* and the ABA-responsive genes *RD29A* and *AREB2*. In addition, SA antagonized ABA’s ability to suppress seed germination. The combined observations that i) SA-deficient *sid2-1* seeds germinated more slowly than WT seeds, and ii) *sid2-1* seeds were hypersensitive to exogenously supplied ABA, argue that endogenous SA plays an important role in counteracting the effects of both endogenously and exogenously supplied ABA.

To investigate the mechanism through which SA antagonizes ABA signaling, we monitored the interaction between several ABA receptors and PP2Cs. SPR analyses revealed that the clade A PP2Cs, ABI1 and ABI2, bind PYL1, PYL2, and PYR1 even in the absence of ABA (Figure 7A); however, these interactions were strongly enhanced in the presence of ABA (Figure 7B). Strikingly, SA suppressed the ABA-enhanced interaction between these proteins, albeit to varying extents depending on the identity of the interacting partners (Figure 7C). Consistent with these results, both *in vitro* and *in vivo* analyses have previously demonstrated that ABA strongly enhances binding between clade A PP2Cs and certain ABA receptors, including PYL1, PYL2 and PYR1 (Park *et al*., 2009). Crystal structure analyses further demonstrated that this interaction inhibits PP2C activity by occluding PP2C’s active site (Melcher *et al*., 2009; Miyazono *et al*., 2009). Since PP2Cs repress ABA signaling by preventing autophosphorylation-dependent activation of SnRK2s (Umezawa *et al*., 2009; Soon *et al*., 2012), ABA-induced binding of PP2Cs by ABA receptors is a critical step in activating ABA signaling (Fujii *et al*., 2009). SA’s ability to suppress the interaction between ABI1 or ABI2 and the ABA receptors therefore provides one mechanism through which SA can antagonize ABA signaling. In addition, our cell-free degradation assay revealed that SA suppresses the ABA-enhanced turnover of PP2Cs and stabilization of SnRK2s. Given that ABI1 was degraded substantially more rapidly in extracts from *sid2-1* mutants than from WT plants, endogenous SA appears to play an important role in regulating cellular PP2C levels. Taken together, these results suggest that SA antagonizes ABA signaling via multiple mechanisms that both promote the enzymatic activity and/or protein stability of negative regulators and decrease the stability of downstream effectors.

**Figure 7:**
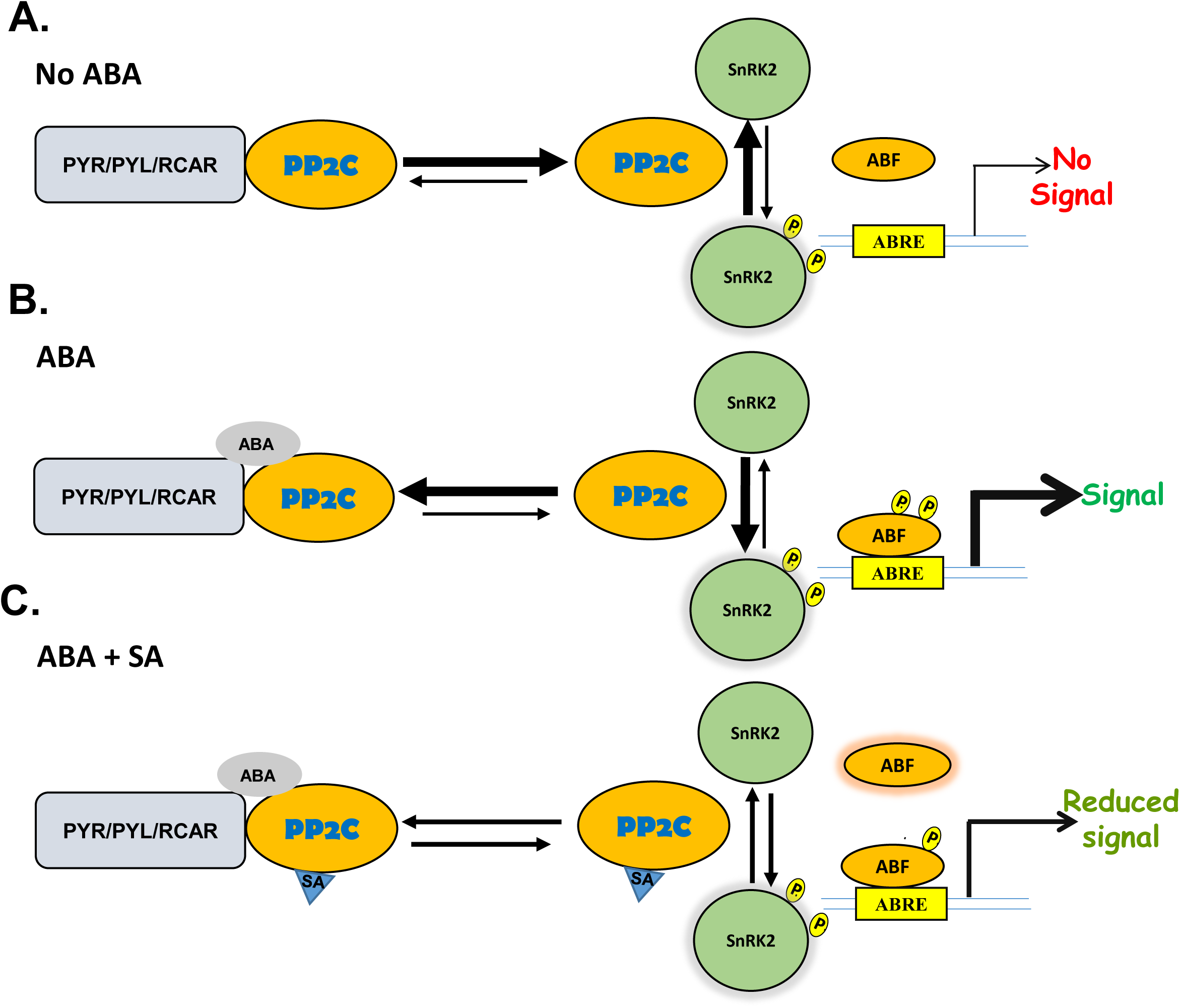
Schematic illustrating part of SA’s antagonistic effects on the ABA signaling module. **(A).** In the absence of ABA, free PP2Cs prevents autophosphorylation-dependent activation of SnRK2s by dephosphorylating them. **(B)** In the presence of ABA, PYR/PYL receptors tightly bind to PP2Cs, thereby preventing free PP2C-mediated dephosphorylation of SnRK2s. Receptor-mediated occlusion of PP2Cs allows autophosphorylated-activated SnRK2s to relay the ABA signaling by phosphorylating downstream targets such as abscisic acid responsive elements-binding factor 2 (ABF2), which enables its binding to ABA-responsive elements (ABRE) in the promoter region of ABA-responsive genes. **(C)** SA suppresses ABA’s enhancement of the interaction of PP2Cs with the PYR/PYL/RCAR receptor and the resulting autophosphorylation-dependent activation of SnRK2s, which results in reduced ABA signaling. The length and thickness of the arrows indicate the equilibrium between free and receptor-bound PP2Cs and between inactive, nonphosphorylated and active, autophosphorylated SnRK2s.

Within this overall framework, differences among the binding specificities and affinities, protein-protein interactions and/or stability of various ABA signaling components may further influence SA/ABA crosstalk. For example, while ABI1 and ABI2 bound SA, another clade A member, HAB1 did not. ABI1 differs from ABI2 as it displayed substantially greater affinity for all three ABA receptors in the absence of ABA; it also was the most stable PP2C in our *in vitro* degradation assay. SA’s ability to disrupt the ABA-enhanced interactions between ABA receptors and PP2Cs also varied, depending on the proteins involved. In particular, the interaction between PYR1 and ABI1 or ABI2 was suppressed less effectively by SA than the interactions between these PP2Cs and the other ABA receptors. Similar to these findings, reconstitution of the ABA signaling pathway in Arabidopsis protoplasts using different combinations of ABA receptors, PP2Cs and SnRK2s previously revealed that the intensity of interactions varied significantly depending on which members of the protein families were involved (Fujii *et al*., 2009). Combined with our findings, these results suggest that while the various the members of the PP2C, ABA receptor and SnRK2 families serve overlapping functions, differences in their temporal and/or spatial expression patterns, as well as their affinity for specific interacting partners and/or SA and ABA, could fine-tune ABA signaling and regulate crosstalk with the SA pathway.

In comparison to the clade A PP2Cs, members of clade D were recently shown to negatively regulate cell expansion by dephosphorylating, and thereby inactivating plasma membrane H^+^-ATPases (Spartz *et al*., 2014). In the presence of auxin, this suppression is relieved by members of the SAUR (small auxin up-regulated) protein family, which bind and inhibit PP2C-Ds. Interestingly, while different SAURs inhibited the activity of several PP2C-D family members (including PP2C-D4) to varying extents, they did not inhibit the activity of a clade A PP2Cs, ABI1, a clade E member, or a phosphatase belonging to the PPP family. Our studies provide additional insights into PP2C-D function, as we demonstrate that PP2C-D4 is an SABP, whose binding to 3AESA is enhanced by ABA. PP2C-D4 also bound several ABA receptors, and this interaction was enhanced by the presence of ABA. In comparison to the clade A PP2C-ABA receptor interactions, however, the binding between PP2C-D4 and the ABA receptors was substantially weaker and its suppression by SA was less effective. In addition, SA consistently stimulated this interaction in the absence of ABA. The expression pattern of *PP2C-D4* in the presence of ABA and/or SA also differed significantly from that of the clade A PP2Cs. Thus, while PP2C-D4 and the clade A PP2Cs share some common features, their many differences are consistent with the previous demonstration that clade D and clade A PP2Cs serve distinct functions.

In response to stress, plants maximize their chances of survival and reproduction by redistributing cellular resources from growth and developmental processes to defensive responses (Asselbergh *et al*., 2008; Atkinson & Urwin, 2012). Many studies have assessed plant responses to individual stresses, but there is a growing recognition that plants in the field contend with multiple stresses simultaneously, and that, depending on the specific stresses, the responses may be additive or antagonistic. Thus, the hormones responsible for mediating developmental processes and stress responses are involved in complex crosstalk that ultimately allows the plant to tailor its response to the environmental conditions. A previous study demonstrated that ABA can interfere with SA-mediated innate immune responses by down-regulating SA biosynthesis (De Torres Zabala *et al*., 2009). By contrast, our results provide a novel mechanism through which SA can antagonize ABA by interfering with multiple aspects of the ABA signaling pathway. The discovery that PP2C-D4 binds SA and several ABA receptors, and that this binding is enhanced in the presence of ABA, suggests an additional mechanism through which SA and ABA can negatively regulate auxin-mediated growth and developmental processes. Indeed, ABA was shown to suppress hypocotyl elongation, and this correlated with dephosphoryloation of H^+^-ATPases (Hayashi *et al*., 2014). These ABA-induced responses were suppressed in the *abi1-1* mutant, suggesting that clade A PP2Cs are involved in auxin-mediated physiological processes. Future studies will be required to determine whether PP2C-D4 and/or other clade D PP2Cs also mediate ABA antagonism of cell expansion, and whether SA binding by PP2C-D4 affects its ability to interact with SAURs and thereby impact auxin signaling.

In summary, we have elucidated SA mechanisms of action in negatively regulating ABA signaling, which likely serves to properly balance the plant response to multiple stresses. As plants confront a changing climate, this balance and our understanding of how it is regulated and might be beneficially altered, take on increase significance.

## Materials and Methods

### Plant materials and growth conditions

The wild-type and *sid2-1 Arabidopsis thaliana* ecotype Col-0 plants were grown on standard Murashige and Skoog (MS) media containing half-strength of MS with pH adjusted to 6.0 by KOH and supplemented with 10 g /L sucrose. Arabidopsis seeds were first surface-sterilized by soaking in a solution of 30% bleach with 0.1% triton X-100 for 5-10 min and then rinsed five times with sterile water. The surface sterilized seeds were incubated at 4 ^°^C for 2 days for stratification before planting on the MS media. For the seed germination assay, (±) abscisic acid (Caisson labs) or salicylic acid (Sigma) were added directly into the MS media. The plates with seed were placed vertically in the growth chamber with 16/8h light/dark cycle, 22 ^°^C, and 70% humidity. The germination rate was measured. For spray treatment, one-week old seedlings were subjected to water, ABA, SA, or ABA+SA spray treatment and whole seedlings were collected for RNA analysis. For cell-free degradation assay, ten-day-old wild-type or *sid2-1* seedlings were subjected to water, ABA, SA, or ABA+SA spray treatment to compare the effects of protein extracts on the stability of ABI1.

### Cloning and plasmid constructs

All oligonucleotides used for cloning and plasmid construction are listed in Table S1. ORFs of PP2CD, ABI1, ABI2, and HAB1were amplified from an Arabidopsis cDNA library. The resulting PCR products were digested with *NdeI* and *BamHI* for ABI1, *NdeI* and *SacI* for ABI2, and HAB1 and cloned into the expression vector pET28a (EMD Millipore, MA, USA) for expression. PP2CD was cloned into pET42a (EMD Millipore, MA, USA) using *NdeI* and *XhoI* cloning sites. Cloning of PYL1, PYL2, PYR1, SnRK2.2, SnRK2.3, and SnRK2.6 into pSUMO- H6SUMO vector were described previously (Soon *et al*., 2012).

### Protein purifications

Two-step protein purifications were performed as described previously (Manohar *et al*., 2017). Briefly, the Rosetta 2 (DE3) (EMD, Millipore, MA, USA) bacterial cells were grown at 37 ^°^C in LB medium containing 50 μg/ml kanamycin and 34 μg/ml chloramphenicol until the OD_600_ of the culture reached approximately 0.6 before addition of isopropyl-β-D-thiogalactoside (IPTG) to a final concentration of 1 mM to induce expression. Induced culture was incubated overnight at 20°C. The cells were then harvested by centrifugation and the pellet was resuspended in the lysis buffer (50 mM Tris pH 7.5, 500 mM NaCl, 10% glycerol, 20 mM imidazole, 0.5% triton X-100 and 1 mM phenylmethylsulphonyl fluoride) and disrupted by sonication. The clarified supernatant obtained after centrifugation was incubated with Ni-NTA His resin (Novagen, MA, USA) and the bound protein was eluted in lysis buffer supplemented with 250 mM imidazole. The eluted proteins were then subjected to gel filtration chromatography on a HiLoad 16/600 Superdex 200 prep grade column (GE Healthcare, PA, USA), using gel filtration buffer (50 mM Tris pH 7.5, 150 mM NaCl, and 10% glycerol). The two-step purified proteins were stored at −80 °C.

### Assessment of 3AESA-binding activities by Surface Plasmon Resonance (SPR)

SPR analyses of 3AESA binding and competition by SA were performed with a Biacore 3000 instrument (GE Healthcare) as described previously (Manohar *et al*., 2015). Immobilization of 3AESA on the CM5 sensor chip was performed as described previously (Tian *et al*., 2012). To test SA-binding activity, proteins were diluted in HBS-EP buffer (GE Healthcare) and passed over the sensor surface of the 3AESA-immobilized and mock-coupled flow cells. The specific binding signal was determined by subtracting the signal generated with the mock-coupled flow cell from the signal generated by the 3AESA-immobilized cell. To re-use the sensor chips, bound proteins were stripped off by injecting NaOH solution (pH 12).

### Assessment of protein-protein interactions by SPR

Protein interaction analyses were performed by SPR using Biacore 3000 instrument. His-SUMO-tagged PYL1, PYL2, and PYR1 were immobilized on a CM5 sensor chip by amine coupling (GE healthcare), essentially by following the manufacturer’s instructions. Briefly, proteins were diluted in 10 mM sodium acetate, pH 5.0 buffer at a concentration of to 50μg/ml. CM5 sensor chip surface was activated by injecting 85 μl of EDC/NHS solution with a flow rate of 10 μl/min. After activation, protein solution was injected for 42 minutes with a flow rate of 10 μl/min. Finally, 85 μl of ethanolamine was flowed over the surface to deactivate remaining active groups and remove non-covalently bound protein with a flow rate of 10 μl/min. The protein immobilization level was stabilized for 12 hours by flowing HBS-EP buffer with a flow rate of 10 μl/min. To test protein-protein interactions, protein analytes (PP2Cs and SnRK2s) were diluted to desirable concentration in HBS-EP buffer in the presence or absence of various concentrations of ABA, SA, or ABA+SA, and then passed over the protein-immobilized sensor surface and mock coupled flow cells with a flow rate of 30 μl/min. The higher flow rate was used to avoid mass-transfer as recommended by the manufacturer. The binding signal was generated by subtracting the signal for mock-coupled flow cells from that for the protein-immobilized flow cells. To re-use the chip, bound proteins were stripped off by injecting 8 μl of 10 mM glycine-HCl solution (pH 3) with a flow rate of 30 μl/min.

### RNA analyses

Unless stated otherwise, at least three biological replicates were used for all RNA analyses. For each replicate, total RNA from one-week old Arabidopsis seedlings was isolated from a pool of five seedlings. Total RNA was isolated from using Qiagen RNeasy plant mini kit (Qiagen) according to the manufacturer instructions. DNAse treatment was done using DNA-free kit (Ambion) following the manufacturer’s instructions. First strand cDNA was synthesized from 1 mg of RNA using M-MLV reverse transcriptase (Promega). For quantitative real-time PCR, transcripts were amplified using SYBR premix Ex Taq II (Takara) with gene-specific primers listed in table S1. Reactions were done using a CFX96 touch Biorad Real-time PCR system (Bio-Rad). The PCR conditions were 95 ^°^C for 3 min (initial denaturation) followed by 44 cycles of amplifications (95 ^°^C for 10s, 60 ^°^C for 30s), followed by generation of a dissociation curve. The relative fold change was calculated according to the 2^-▲ ▲ Ct^ method (Manosalva *et al*., 2015). Ubiquitin was used as endogenous reference gene. The paired t-test with an α-level of 0.05 was used to compare transcript level in the ABA, SA, ABA+SA, and mock-treated plant samples.

### Cell-free degradation assay

The tissue samples were collected from 10 day old seedlings of wild-type and *sid2-1* and finely grounded using liquid nitrogen. The total protein extracts were then prepared using protein extraction buffer (25 mM Tris pH 7.5, 10 mM NaCl, 10 mM MgCl_2_, and 4 mM PMSF). The sample was vortexed to mix and centrifuged twice at 17,000 g for 10 min at 4 ^°^C to remove debris. The clear supernatant was pre-treated with 1 mM cycloheximide (MP Biomedicals) for 1 h to inhibit *de novo* protein biosynthesis. The extracts were then adjusted to equal protein concentrations in degradation buffer (25 mM Tris pH 7.5, 10 mM NaCl, 10 mM MgCl_2_, and 4 mM PMSF, 5 mM DTT, and 10 mM ATP). For degradation assay, an equal amount (approximatly 500 ng) of purified PP2Cs, SnRK2s, and PYL1 were incubated in 50 μl of Arabidopsis total protein extract (containing approximatly 100 μg total proteins) at 28 ^°^C for 3 h, unless otherwise indicated. For ABI1 and SnRK2.6 twice as much total protein extract (approximately 200 μg) was used to more clearly visualize the effect of SA (Figure 4A, 4B). Immunoblot analyses were performed to detect protein levels by using an α-His_6_-HRP polyclonal antibody (QED Biosciences).

## Acknowledgement

We thank D’Maris Dempsey for editing and the US National Science Foundation for financial support via grant IOS-0820405 to D.F.K. We thank Prof. Karsten Melcher (Van Andel Research Institute), for the *E. coli* expression vectors corresponding to PYL1, PYL2, PYR1, SnRK2.2, SnRK2.3, and SnRK2.6.

## Authors’ contributions

M.M, P.M, and D.F.K conceived the research. M.M, D.W, E. K, and D.F.K designed the research. M.M, D.W, P.M, and H.W.C performed the research. M.M, D.W, E.K, and D.F.K analyzed the data. M.M and D.F.K wrote the paper. All authors have read and approved the final manuscript.

## Supplementary Figures

**Supplementary Figure 1:**
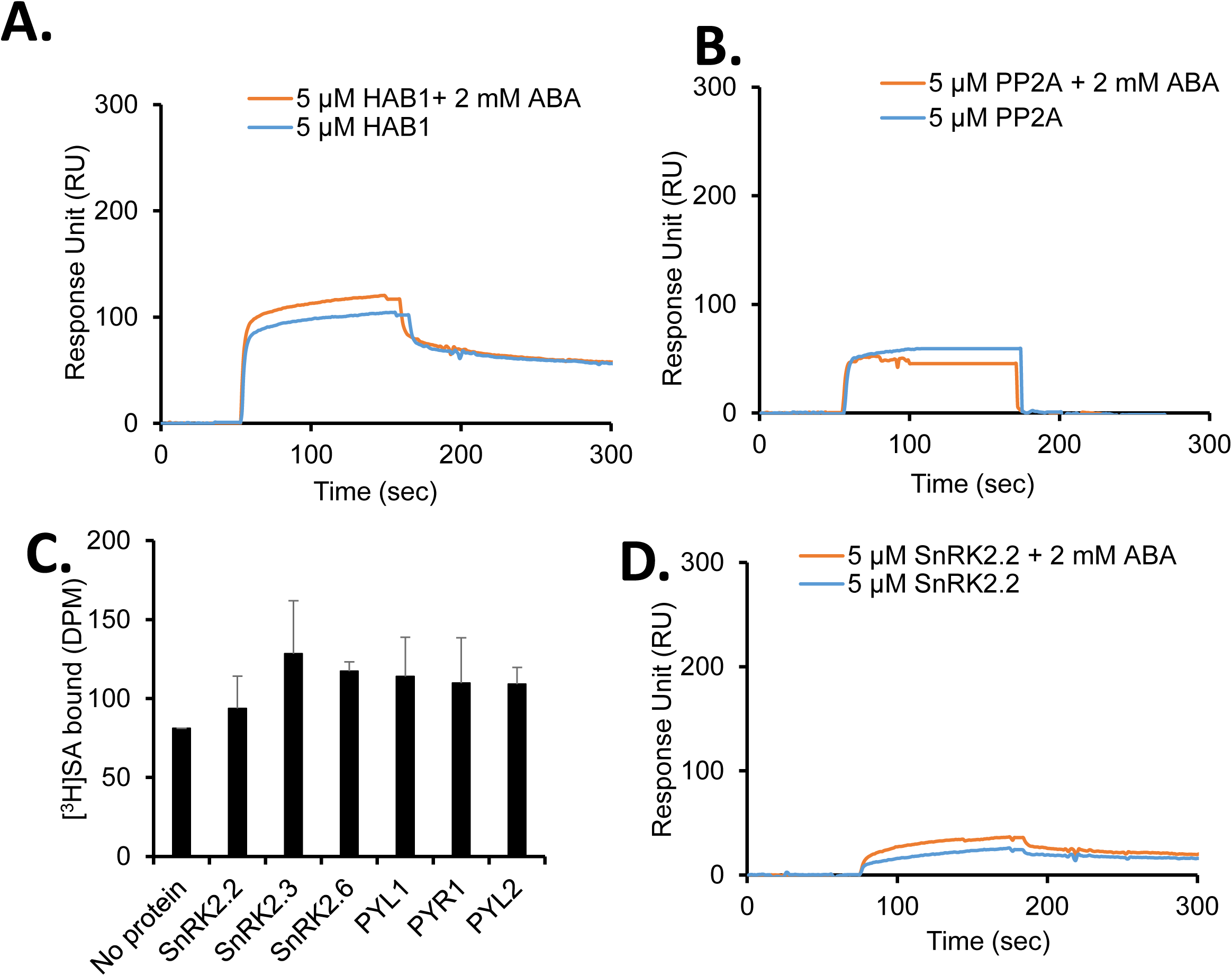
HAB1, PP2A and members of the PYR/PYL/RCAR and SnRK2 families are not SABPs. **(A-C)** Sensorgrams obtained using a 3AESA-immobilized SPR sensor chip with 5 μΜ of His_6_-HAB1 (At1g72770), His_6_-PP2A (At1g25490), or His_6_-SUMO-SnRK2.2 (At3g50500) in the absence or presence of 2 mM ABA. Signals detected from a mock-coupled control chip were subtracted. **(D)** Binding of [^3^H]SA (200 nM) by 200 ng/μl SnRK2.2, 2.3, 2.6, PYL1, PYL2, or PYR1 was determined by size-exclusion chromatography. Chromatography with [^3^H]SA in the absence of protein served as negative control.

**Supplementary Figure 2:**
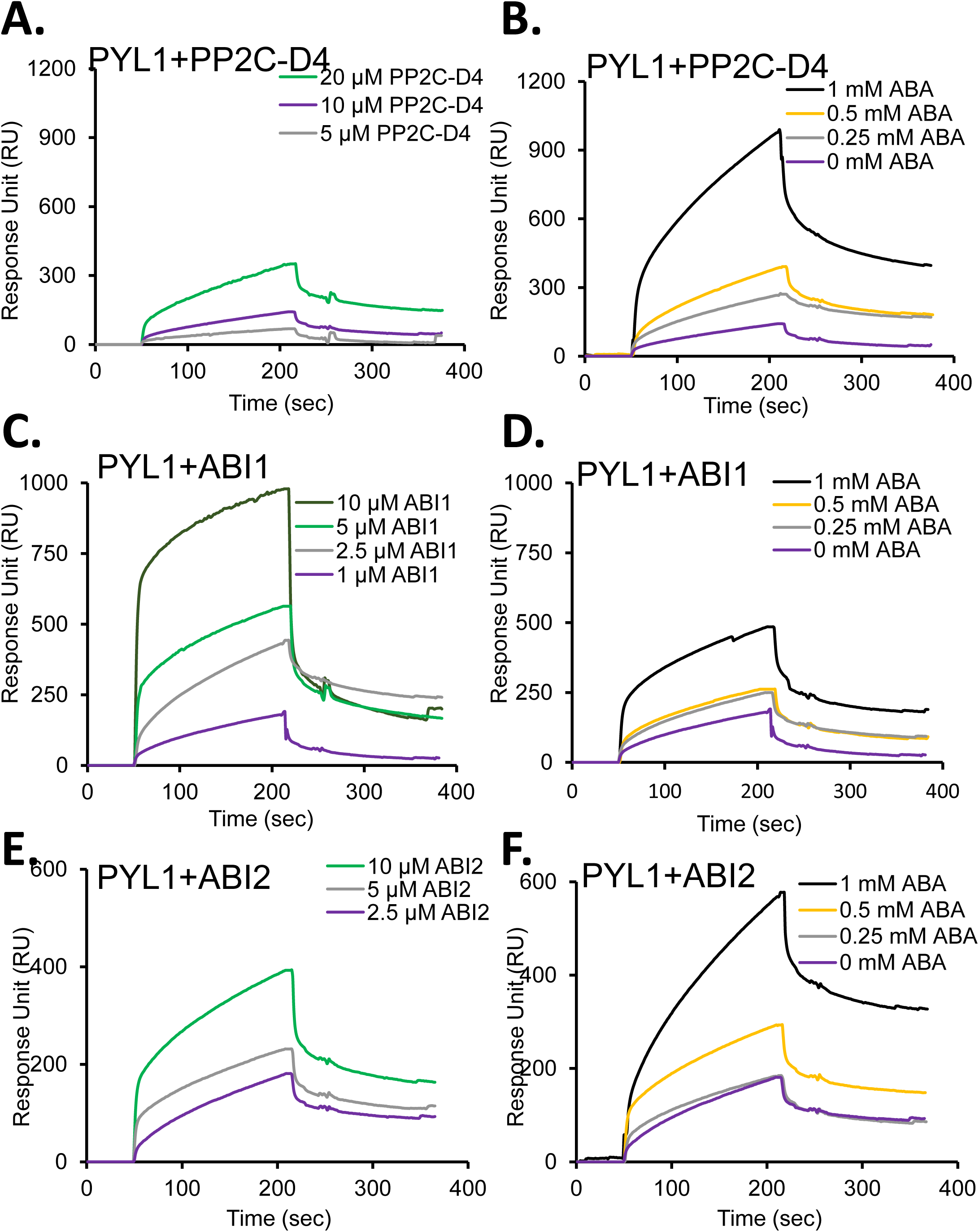
Effect of ABA on the interactions of PP2Cs with PYL1. **(A, C, E)** Sensorgrams obtained with the indicated concentrations of recombinant, purified His_6_- tagged PP2C-D4, ABI1, or ABI2 using a His_6_- SUMO-tagged PYL1-immobilized sensor chip**. (B, D, F)** ABA dose-dependent effect on the interactions between 10 μΜ PP2C-D4, 1 μΜ ABI1, or 2.5 μΜ ABI2 and His_6_- SUMO-tagged PYL1 immobilized on the sensor chip. Signals detected from a mock-coupled control chip were subtracted. The experiments was independently repeated at least twice.

**Supplementary Figure 3:**
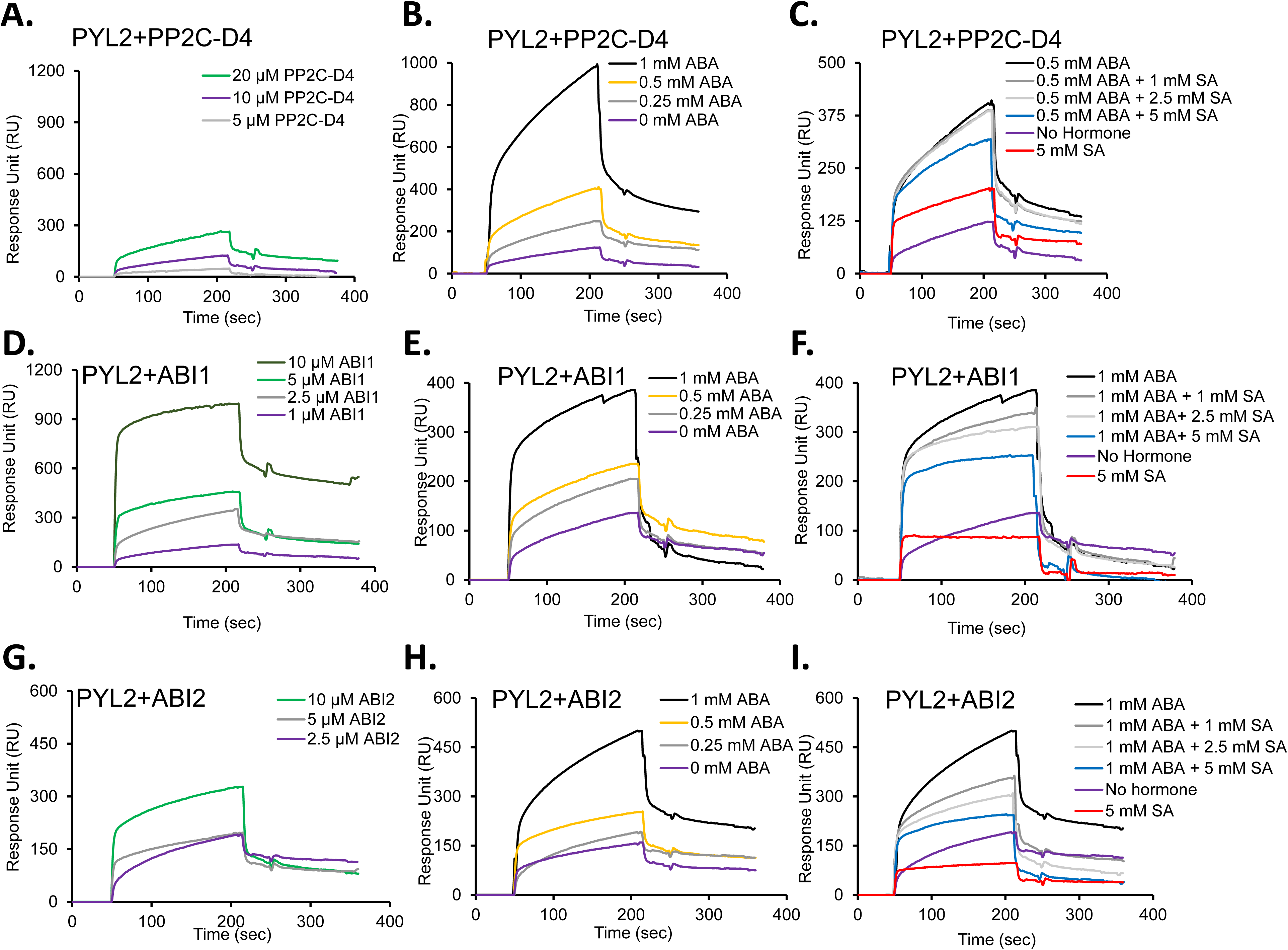
SA disrupts the ABA-enhanced interactions between PP2Cs (PP2C-D4, ABI1, and ABI2) and the ABA receptor PYL2. **(A, D, G)** Sensorgrams obtained using a His_6_-SUMO-tagged PYL2-immobilized sensor chip and the indicated concentrations of recombinant, purified His_6_- tagged PP2C-D4, ABI1, or ABI2**. (B, E, H)** ABA dose-dependent effect on the interactions of 10 μM PP2C-D4, 1 μM ABI1, or 2.5 μM ABI2 with His_6_-SUMO-tagged PYL2-immobilized sensor chip. **(C, F, I)** Sensorgrams obtained with 10 μΜ of the His_6_-tagged PP2C-D4, 1 μM ABI1, or 2.5 μM ABI2 using a His_6_-SUMO-tagged PYL2-immobilized sensor chip in the absence or presence of the indicated concentrations of ABA, SA or ABA plus SA. Signals detected from a mock-coupled control chip were subtracted. The experiments was independently repeated at least twice.

**Supplementary Figure 4:**
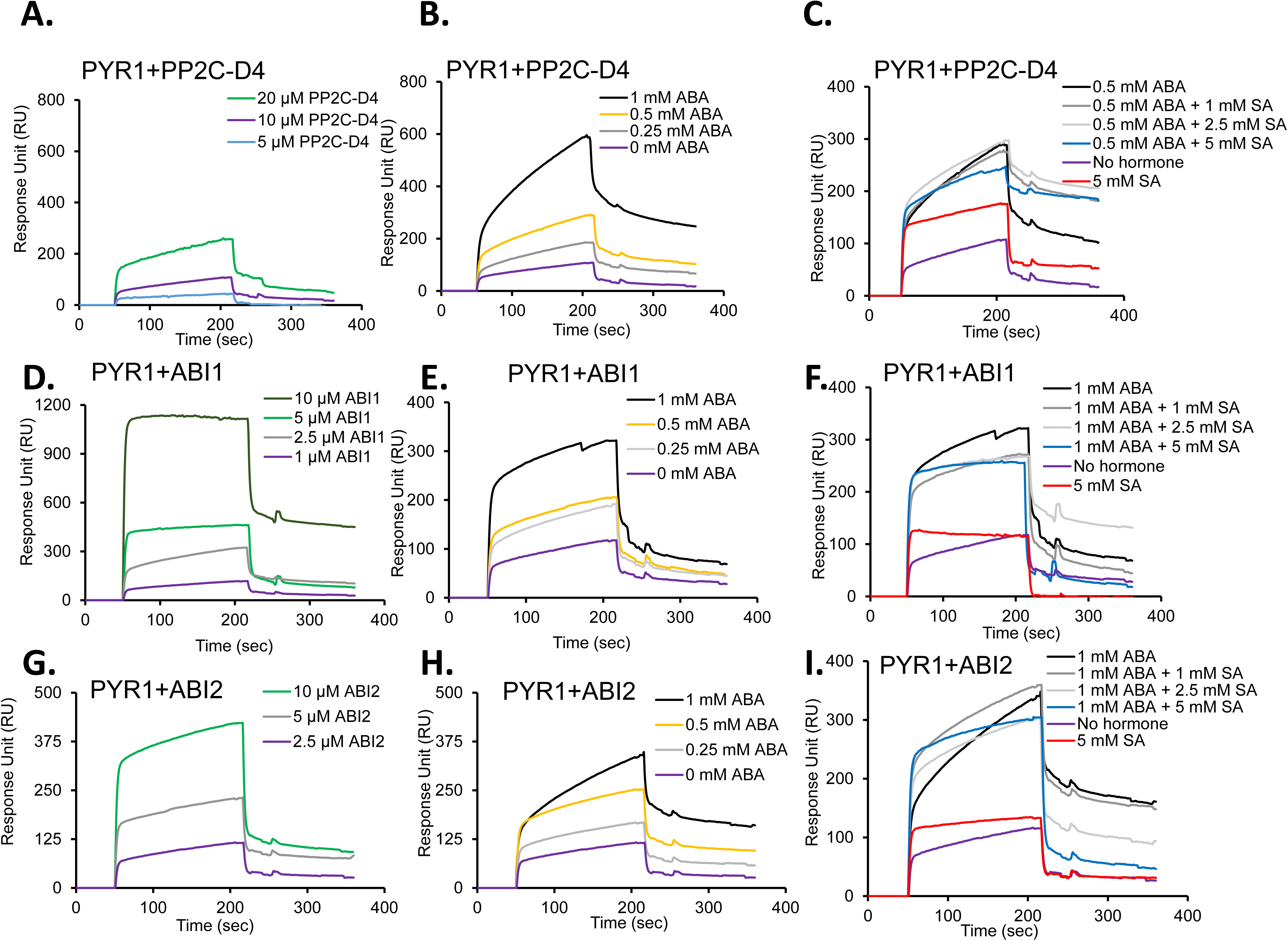
SA disrupts the ABA-enhanced interactions between PP2Cs (PP2C-D4, ABI1, and ABI2) and ABA receptor PYR1. **(A, D, G)** Sensorgrams obtained using a His_6_-SUMO-tagged PYR1-immobilized sensor chip and the indicated concentrations of recombinant, purified His_6_- tagged PP2C-D4, ABI1, or ABI2**. (B, E, H)** ABA dose-dependent effect on the interactions of 10 μM PP2C-D4, 1 μM ABI1, or 2.5 μM ABI2 with His_6_-SUMO-tagged PYR1-immobilized sensor chip. **(C, F, I)** Sensorgrams obtained with recombinant 10 μΜ of the His_6_- tagged PP2C-D4, 1 μM ABI1 or 2.5 μM ABI2 using a His_6_- SUMO-tagged PYR1-immobilized sensor chip in the absence or presence of the indicated concentrations of ABA, SA, or ABA plus SA. Signals detected from a mock-coupled control chip were subtracted. The experiments was independently repeated at least twice.

**Supplementary Figure 5:**
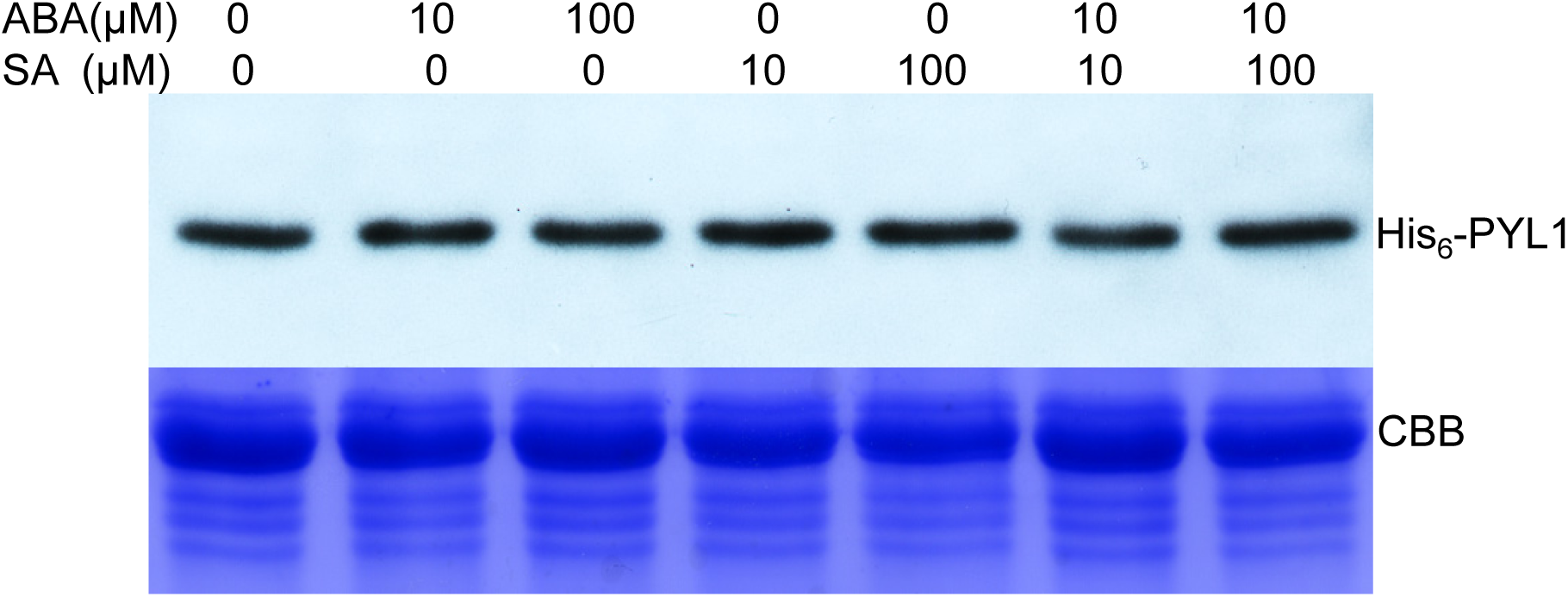
ABA and SA alone or in combination do not affect turnover of PYL1. Cell-free degradation assay using total protein extracts prepared from ten-day-old Arabidopsis seedlings supplemented with 500 ng of His_6_-Sumo-tagged PYL1 and indicated concentrations of ABA, SA, or ABA+SA. The degradation assay was carried out at 30^0^ C for 3 hrs. Proteins were detected by immunoblotting using an α-His_6_-HRP antibody and Coomassie brilliant blue (CBB) staining of the gel served as a loading control. The experiments was independently repeated twice.

**Supplementary Figure 6:**
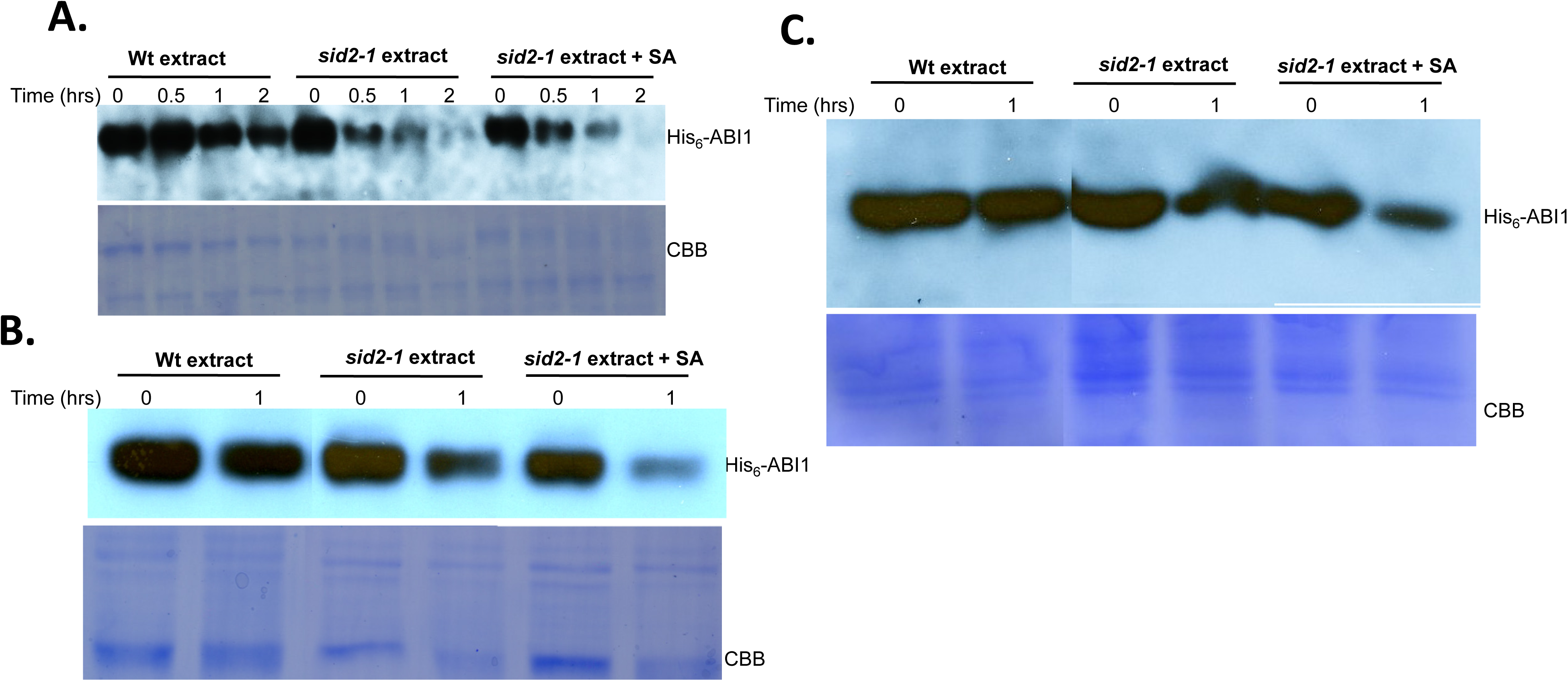
SA does not affect the rate of ABI1 degradation in *sid2-1*plants. **(A)** Cell-free degradation assay using 500 ng of His_6_- tagged ABI1 and Arabidopsis protein extracts from Wt or *sid2-1* mutant plants in the absence or presence of 10 μΜ SA that was added directly to the reaction mix. His_6_- tagged ABI1 was detected in samples harvested at the indicated times by immunoblotting using a α-His_6_-HRP antibody. **(B)** Cell-free degradation assay using 500 ng of His_6_-tagged ABI1 and Arabidopsis protein extracts from ten-day-old Wt or *sid2-1* mutant plants that were sprayed with water or 10 μΜ SA three hours prior to extract preparation. His_6_- tagged ABI1 was detected in samples harvested at the indicated times by immunoblotting using a α-His_6_-HRP antibody. **(C)** Cell-free degradation assay using Arabidopsis protein extracts from ten-day-old wild-type (Wt) or SA-deficient *sid2-1* mutant plants grown in MS medium in the absence or presence of 10 μΜ SA and supplemented with 500 ng of His_6_-tagged ABI1. His_6_- tagged ABI1 was detected in samples harvested at the indicated times by immunoblotting using an α-His_6_-HRP antibody. For B & C, all lanes are from the same experiment; some lanes unrelated to this study were removed and lanes were then merged for clarity of presentation. The experiments was independently repeated twice.

